# Moment-to-Moment Pain Variability Predicts Pain Chronification

**DOI:** 10.64898/2026.03.02.709010

**Authors:** Gaia Pantaleo, Carl Ashworth, Maeghal Jain, Flavia Mancini

## Abstract

Moment-to-moment fluctuations in perceived pain are often dismissed as noise. However, across neural systems, rapid variability reflects fundamental network properties such as stability and flexibility, and in pain, reduced variability has been linked to chronic pain severity. Here, we tested whether alterations in such variability mark an early mechanistic transition from subacute to chronic pain. Using a longitudinal dataset of 120 individuals with subacute back pain followed over one year, we analysed continuous pain ratings alongside fMRI activity during spontaneous pain.

We show that patients who develop persistent pain exhibit a marked decrease in moment-to-moment pain variability over the course of one year, in contrast to recovering individuals whose variability increases over time. Neural activity in thalamo–cortico–limbic and modulatory circuits associated with these fluctuations at pain onset predicts clinical outcomes one year later. Notably, similar predictive patterns are observed during the early post-onset, indicating that these neural signatures emerge early and remain stable during the initial phase of disease progression.

These findings indicate that early loss of dynamical flexibility is a hallmark of pain chronification and identify moment-to-moment variability as a clinically accessible marker of the system’s dynamical state.

## Main

Pain fluctuates over time, with moments of relief and moments of heightened intensity. Although these rapid fluctuations are often dismissed as noise, extensive work across neural systems shows that moment-to-moment variability reflects core properties of network dynamics, including stability and flexibility [18, 35, 17, 25, 14, 49, 31, 56, 32]. Variability is therefore increasingly viewed as a fundamental signature of system function rather than measurement error.

Despite this, such short-term variability has received comparatively little attention in pain research. Most prior work has examined fluctuations over days or weeks [87, 70, 37, 30, 39, 63, 38, 13], or has focused on task-evoked pain responses [90, 54]. As a result, the field lacks mechanistic accounts of how rapid fluctuations in perceived pain relate to the early transition from subacute to chronic pain. Yet emerging evidence suggests that reduced moment-to-moment variability accompanies higher pain severity and chronic pain states [91]. This suggests that early alterations in variability reflect a loss of dynamical flexibility within pain-regulatory circuits.

This gap motivates the central question of the present study: Can moment-to-moment variability in spontaneous pain, measured early after pain onset, provide an early marker of who will recover and who will develop persistent pain?

Addressing this question requires a mechanistic understanding of where changes in variability originate. One possibility is that they reflect central reorganisation within the endogenous pain-control system, in which disrupted flexibility across brain, brainstem, and spinal circuits reduces the system’s capacity to transition out of high-pain states [9, 44, 65, 69, 10, 45]. Another possibility is that variability arises from peripheral alterations—such as changes in small-fiber function or aberrant spontaneous activity [72, 24, 82, 64, 33, 43]—that propagate centrally and constrain the global dynamics of the pain-regulatory network [71]. Identifying early neural correlates of variability could therefore illuminate the mechanisms underlying pain chronification while offering a clinically tractable marker.

Existing efforts to identify pain biomarkers have spanned molecular [79, 76, 86, 19, 67, 52, 66], genetic [12], microbiome [77, 55, 5], and neural markers [84, 68, 85, 74, 36, 20, 81, 75, 80, 11, 26]. Notably, recent effort effectively built a corticospinal signature capable of capturing interindividual pain sensitivity [50]. However, most neural markers do not incorporate moment-to-moment variability, instead focusing on longer timescales [48] or experimentally induced pain [88], which do not fully capture the spontaneous temporal dynamics of pain.

A recent study has shown that moment-to-moment fluctuations in spontaneous pain can be decoded from brain activity within individuals, enabling prediction of ongoing pain intensity but not generalisation across individuals [46]. Such approaches focus on tracking current pain states and do not address whether these fluctuations reflect the underlying system state that governs the future clinical trajectory of pain. We propose that moment-to-moment variability provides a direct and measurable window into the dynamical state of pain-regulatory systems and can therefore predict the transition from subacute to chronic pain.

Here, we test the hypothesis that early reductions in short-term pain variability reflect a loss of dynamical flexibility in pain-regulatory networks and predict long-term clinical outcomes. Using a longitudinal cohort of 120 individuals with subacute back pain (SBP; [7, 2]) followed over one year (fig. 1A), we ask: (1) Does moment-to-moment variability evolve differently in individuals who recover versus those who develop persistent pain? (2) Do neural signatures associated with variability during the early post-onset period (first 1–2 months) predict clinical outcome one year later?

**Figure 1.**
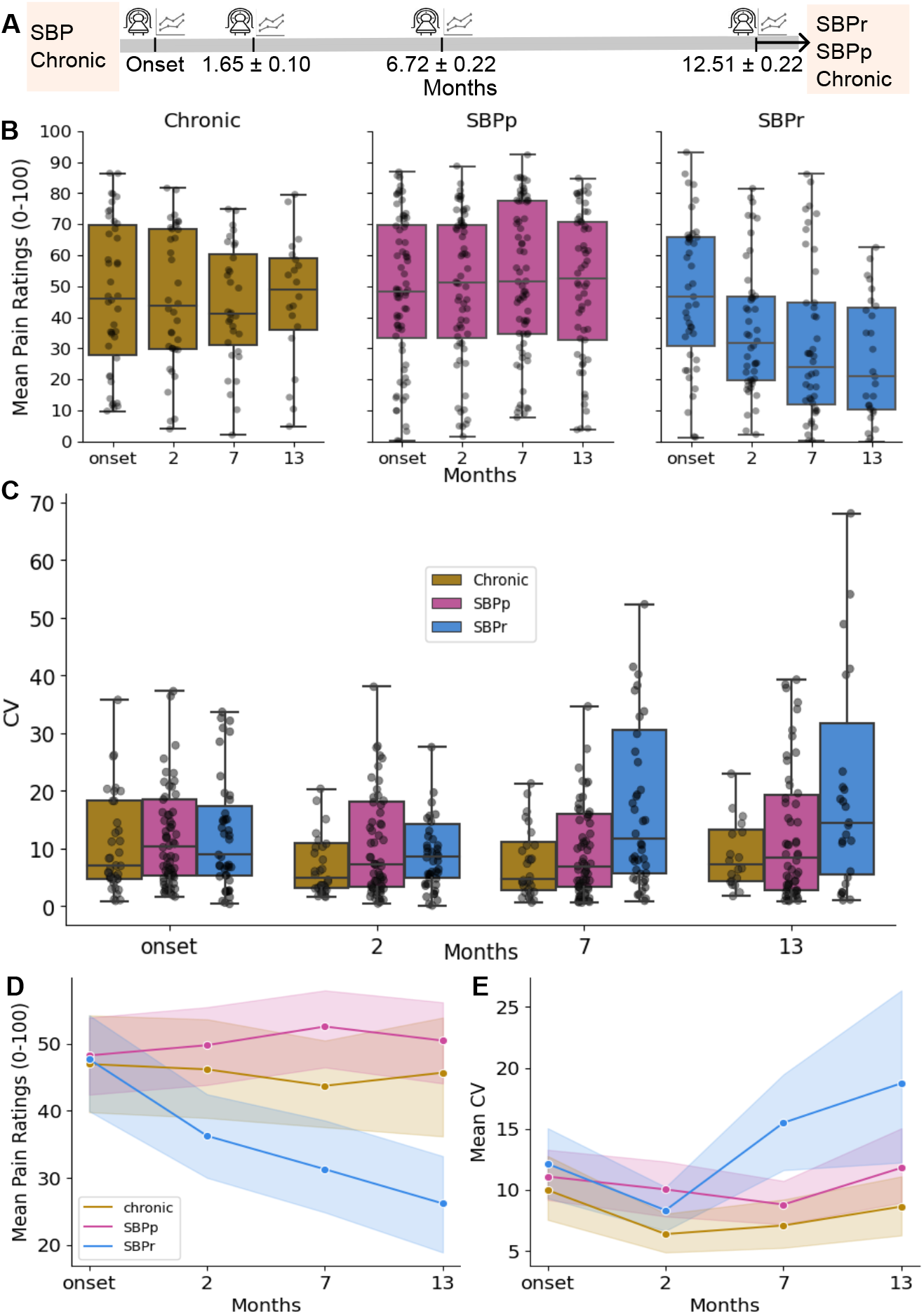
**A.** Study design: 120 participants entered a longitudinal study run over the course of one year by Baliki et al. [7]. The dataset consists of 70 SBP participants, a group of 26 chronic patients and a group of 26 healthy participants. Participants were followed across four visits (at ~ 0, 1.6, 6.7, and 12.5 months), with fMRI scans and continuous pain ratings collected at each. In our analysis we only kept SBP participants and chronic patients, which served as control group. Of the original set of SBP and chronic participants, we kept 93 participants (69 SBP, 24 chronic) after participant selection based on completeness of ratings and fMRI data, following data cleaning. 92 participants were selected in visit 1 (42 SBPp, 26 SBPr, 24 chronic), 83 in visit 2 (40 SBPp, 23 SBPr, 20 chronic), 88 in visit 3 (43 SBPp, 24 SBPr, 21 chronic) and 76 in visit 4 (40 SBPp, 17 SBPr, 19 chronic). **B**. Participants were classified into SBPr or SBPp based on mean pain reduction (≥20% for SBPr); participants formally diagnosed as chronic formed a third group, which was treated as control. **C**. Mean pain decreased in SBPr but remained stable in SBPp and chronic groups. **D**. SBPr Participants showed increasing inter-subject variability in pain (CV) over time, unlike SBPp and chronic groups. **E**. Mean CV rose in SBPr but remained stable in SBPp and chronic groups.

## Results

### Pain variability differentiates longitudinal clinical trajectories

We analysed a longitudinal cohort of individuals with subacute back pain followed over one year, combining continuous moment-to-moment pain ratings with fMRI recordings acquired across four visits (Fig. 1A; [7]). Participants were classified based on clinical outcome into recovering (SBPr), persistent (SBPp), and chronic groups (see Methods). After applying inclusion criteria, 63 subacute participants were retained in visit 1 and 23 chronic patients (SBPr = 26, SBPp = 42). 55 participants completed the study at visit 4 (SBPr = 17, SBPp=40) and 19 chronic patients.

We first tested whether variability in pain ratings distinguishes longitudinal clinical trajectories by examining how the coefficient of variation (CV) of pain ratings evolves over time. An Aligned Rank Transform (ART) ANOVA revealed significant main effects of group (*F*_2,452_ = 10.43, *p <*.001), visit (*F*_3,452_ = 7.00, *p <*.001), and a significant interaction between group and visit (*F*_6,452_ = 3.75, *p* = 0.0012). In particular, the SBPr group exhibited a marked increase in CV across sessions, indicating increasing pain variability over time, with a marked increase in the last session. In contrast, SBPp and chronic volunteers exhibited stable CV values (Figure 1D). ART-C [22] pre-planned contrast tests confirmed divergence of groups in time (omnibus contrast: *p <*.001), demonstrating that despite similar CV magnitudes at individual visits, groups exhibit distinct temporal profiles of variability.

### Neural activity linked to variability predicts clinical outcome

We next tested whether neural activity associated with pain variability predicts clinical outcome, i.e., the risk of developing persistent pain one year later. Two prediction settings were examined: (i) using baseline data at pain onset (visit 1), and (ii) using data from the early post-onset period (visit 2; 1.65 ± 0.10 months). Classification was performed using Gradient Boosting Machine (GBM) at the level of individual regions of interest (ROIs)

Across both prediction settings, the classifier achieved strong performance (Figs. 2, 3). ROC AUC values were consistently above chance, with several ROIs exceeding 0.8. Confusion matrices showed clear diagonal structure, indicating accurate separation of recovering (SBPr) from persistent (SBPp) and chronic participants, with relatively few misclassifications. These results demonstrate that distributed neural activity associated with moment-to-moment variability contains predictive information about future pain trajectories.

**Figure 2.**
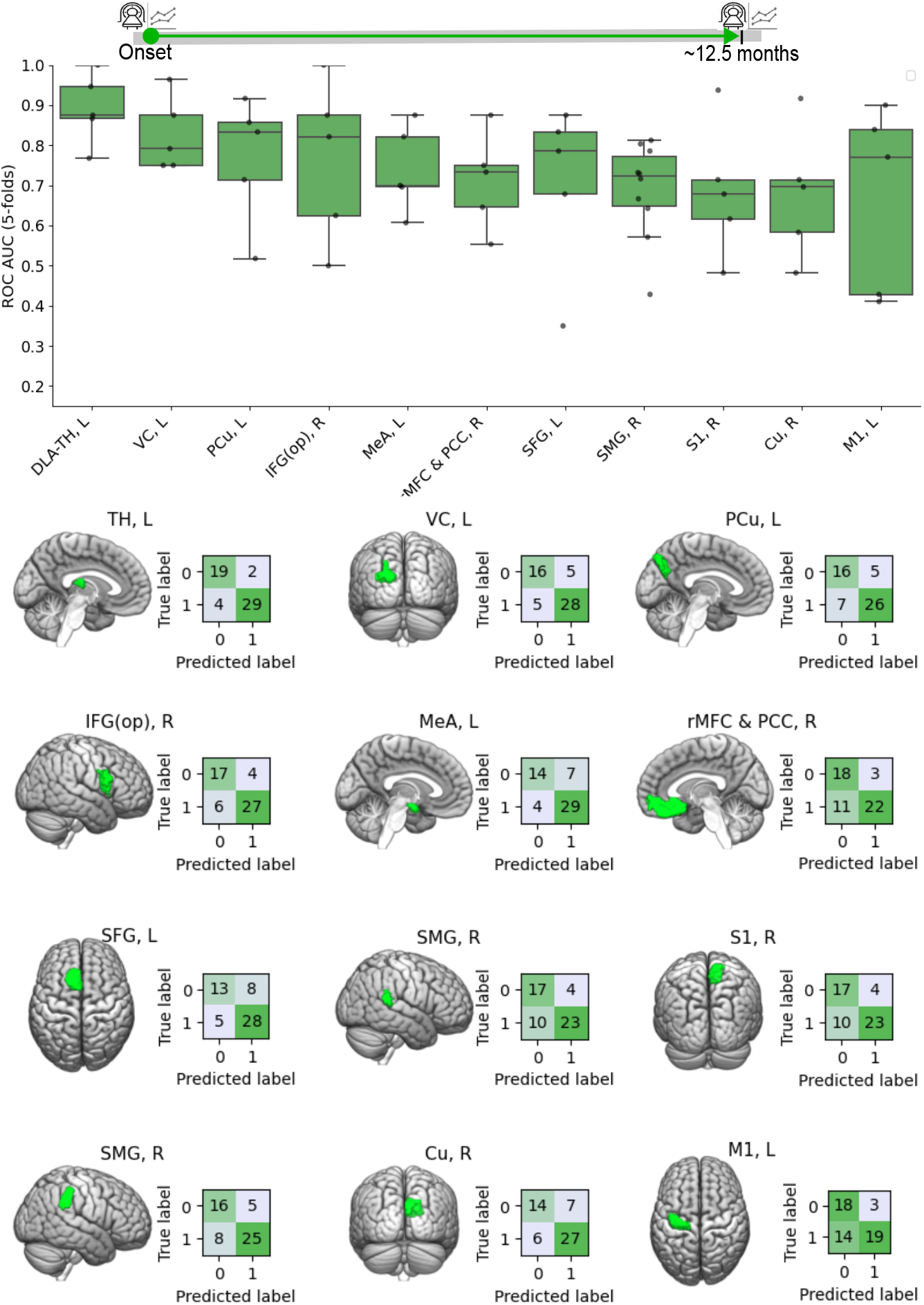
**A.** Top 12 ROIs contributing to classification performance using a Gradient Boosting Machine (GBM) classifier trained on data from the earliest visit to predict clinical outcomes after one year and their associated predictive accuracy. ROI labels correspond to the following anatomical regions: *TH, L* Left Thalamus; *VC, L* Left Visual Cortex; *PCu, L* Left Precuneus Cortex; *IFG(op), R* Right Inferior Frontal Gyrus; *MeA, L* Left Middle Amygdala; *rMFC & PCC, R* Right Rostral Medial Frontal Cortex and Paracingulate Cortex; *SFG, L* Left Superior Frontal Gyrus; *SMG, R* Right Supramarginal Gyrus; *S1, R* Right Primary Somatosensory Cortex; *Cu, R* Right Cuneus Cortex; and *M1, L* Left Primary Motor Cortex.**B**. Cortical surface maps of the corresponding ROIs and their associated confusion matrices at optimal classification threshold.

**Figure 3.**
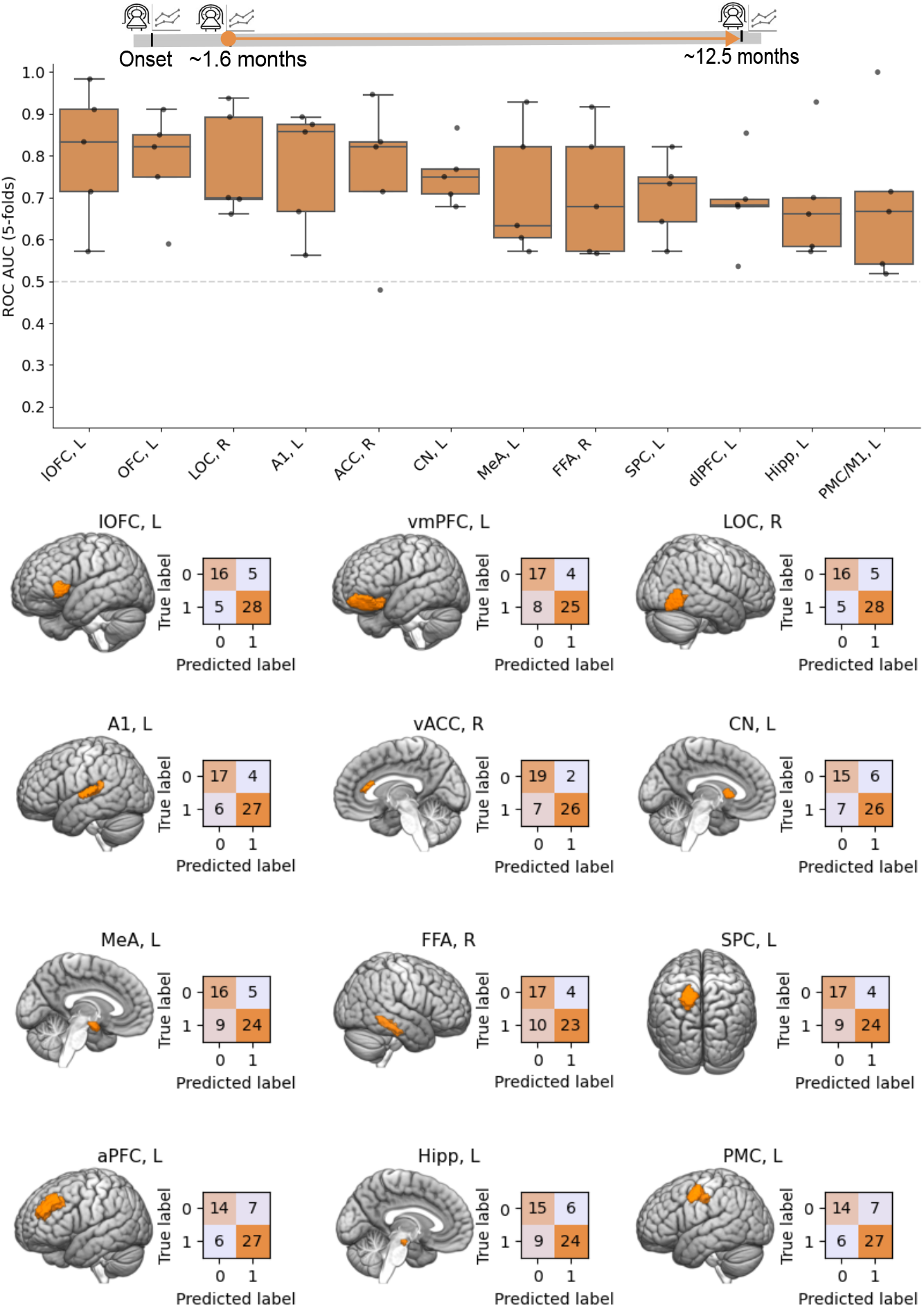
**A.** Top 12 ROIs contributing to classification performance using a Gradient Boosting Machine (GBM) classifier trained on data from ~ 1.65 months post-injury to predict clinical outcomes after one year and their associated predictive accuracy. ROI labels correspond to the following anatomical regions: *lOFC, L* Left Lateral Orbito-Frontal Cortex; *OFC, L* Left Orbito-Frontal Cortex; *LOC, R* Right Lateral Occipital Cortex; *A1, L* Left Primary Auditory Cortex; *ACC, R* Right Anterior Cingulate Cortex; *CN, L* Left Caudate; *MeA, L* Left Middle Amygdala; *PHC, R* Right Parahippocampal Cortex/Lingual Cortex; *SPC, L* Left Superior Parietal Cortex; *dlPFC, L* Left Dorso-Lateral Prefrontal Cortex; and *Hipp, L* Left Hippocampus; and *PMC/M1, L* Left Premotor Cortex/ Motor Cortex. **B**. Cortical surface maps of the corresponding ROIs and their associated confusion matrices at optimal classification threshold.

At pain onset, predictive information was distributed across a set of regions spanning sensory–perceptual, affective, and regulatory components of the pain system (Table 1, Fig. 2). These included thalamic nuclei, prefrontal and frontal regions, as well as the amygdala. Additional contributions arose from regions involved in sensory and perceptual processing, including the primary somatosensory (S1) and motor (M1) cortices, parietal and visual regions.

**Table 1.** MNI coordinates of the top 12 regions whose activity associated with moment-to-moment pain variability at pain onset most strongly predicted clinical outcome at one year. Regions are ranked based on their predictive contribution in the classification model. Values represent mean ROC AUC across 5-fold cross-validation.

| Region | Coordinates (MNI152) |
| --- | --- |
| Left Thalamus (TH) | [-14.65, -19.67, 12.95] |
| Left Visual Cortex (VC) | [-23.09, -87.13, 24.01] |
| Left Precuneus Cortex (PCu) | [-9.53, -73.09, 37.43] |
| Right Inferior Frontal Gyrus, pars opercularis (IFG(op), R) | [51.39, 10.61, 20.25] |
| Left Middle Amygdala (MeA) | [-19.27, -5.27, -16.28] |
| Right Rostral Medial Frontal & Paracingulate Cortex (rMFC & PCC) | [4.91, 36.39, -14.03] |
| Left Superior Frontal Gyrus (SFG) | [-9.31, 16.98, 63.17] |
| Right Supramarginal Gyrus (SMG) | [60.01, -38.60, 16.66] |
| Right Primary Somatosensory Cortex (S1) | [10.18, -39.07, 68.74] |
| Right Supramarginal Gyrus (SMG) | [62.05, -37.48, 37.24] |
| Right Cuneus Cortex (Cu) | [11.30, -73.77, 25.44] |
| Left Primary Motor Cortex (M1) | [-31.30, -19.76, 63.84] |

**Table 2.**
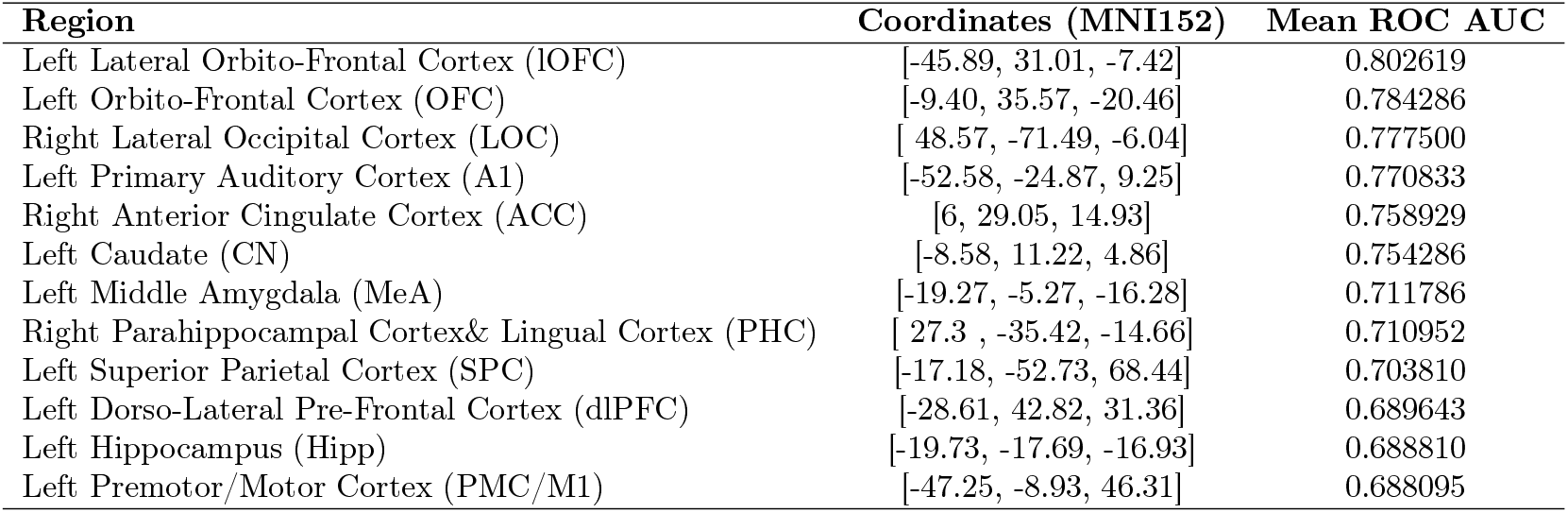
MNI coordinates of the top 12 regions whose activity associated with moment-to-moment pain variability during the early post-onset period (~1.6–1.7 months) most strongly predicted clinical outcome at one year. Regions are ranked based on their predictive contribution in the classification model. Values represent mean ROC AUC across 5-fold cross-validation.

| Region | Coordinates (MNI152) | Mean ROC AUC |
| --- | --- | --- |
| Left Lateral Orbito-Frontal Cortex (lOFC) | [-45.89, 31.01, -7.42] | 0.802619 |
| Left Orbito-Frontal Cortex (OFC) | [-9.40, 35.57, -20.46] | 0.784286 |
| Right Lateral Occipital Cortex (LOC) | [ 48.57, -71.49, -6.04] | 0.777500 |
| Left Primary Auditory Cortex (A1) | [-52.58, -24.87, 9.25] | 0.770833 |
| Right Anterior Cingulate Cortex (ACC) | [6, 29.05, 14.93] | 0.758929 |
| Left Caudate (CN) | [-8.58, 11.22, 4.86] | 0.754286 |
| Left Middle Amygdala (MeA) | [-19.27, -5.27, -16.28] | 0.711786 |
| Right Parahippocampal Cortex & Lingual Cortex (PHC) | [ 27.3 , -35.42, -14.66] | 0.710952 |
| Left Superior Parietal Cortex (SPC) | [-17.18, -52.73, 68.44] | 0.703810 |
| Left Dorso-Lateral Pre-Frontal Cortex (dlPFC) | [-28.61, 42.82, 31.36] | 0.689643 |
| Left Hippocampus (Hipp) | [-19.73, -17.69, -16.93] | 0.688810 |
| Left Premotor/Motor Cortex (PMC/M1) | [-47.25, -8.93, 46.31] | 0.688095 |

A similar pattern was observed when prediction was performed using data from the early post-onset period (visit 2; ~2 months). Predictive regions largely overlapped with those identified at pain onset (Table 1), including thalamic, prefrontal, limbic, and sensory–perceptual areas. The consistency of these regions across timepoints indicates that predictive information is not transient but reflects a stable, distributed network already present at early stages of the condition. Notably, the persistence of both regulatory (prefrontal cortex, thalamus, amygdala) and sensory–perceptual components suggests that variability captures coordinated interactions between top–down and bottom–up processes from the outset of pain.

Together, these findings indicate that neural signatures of variability associated with future pain chronification are established early—within the first weeks following onset—and remain detectable during the initial phase of disease progression.

To further assess whether these predictive signatures reflected future chronic pain trajectories rather than general differences between recovering and non-recovering participants, we performed supplementary analyses considering only the SBPp and chronic groups (Supplementary Figs. 1, 2). Classification performance was reduced relative to analyses including recovering individuals, but predictive information remained detectable at the earliest visit. By the early post-onset period, the classifier showed increasing difficulty distinguishing SBPp from chronic participants, consistent with greater similarity between these groups over time.

## Discussion

Pain chronification remains poorly understood, in part because the field lacks markers that capture how the nervous system transitions from a flexible, recoverable state to a persistent condition. Here, we show that moment-to-moment variability in pain provides such a marker: neural activity associated with variability predicts clinical outcome one year later and distinguishes individuals who recover from those who develop persistent pain.

Recent work has demonstrated that spontaneous pain fluctuations can be decoded from brain activity to track ongoing pain intensity within individuals [46]. In contrast, our results show that variability itself reflects the underlying dynamical state of the pain system and predicts its future clinical trajectory. Notably, these predictive signatures are present at pain onset and remain largely stable during the early post-onset period (~ 1–2 months), indicating that the neural signature of pain chronification is established early rather than progressively emerging.

Together, these findings suggest that variability is not merely a feature of ongoing pain representation, but a readout of the system’s dynamical state—capturing its evolution over time and progression toward chronic pain. In this sense, variability functions as a low-dimensional descriptor of system state, linking moment-to-moment fluctuations in experience to long-term disease progression.

These results align with our previous work [91], which showed that reduced moment-to-moment variability of chronic MSK pain inversely correlates with greater severity. More broadly, they support the view that variability is an inherent property of brain dynamics, reflecting stability and flexibility of a system [15, 4, 6, 27, 41, 58, 59]. These observations suggest that moment-to-moment variability should not be dismissed as noise, but instead interpreted as a window into the system’s ongoing flexibility, providing a simple and informative readout of future clinical trajectories.

### Sources of variability

Although we can identify predictive regions based on changes in their moment-to-moment variability, the origins of these shifts remain unclear. One possibility is that they reflect altered functioning of the endogenous modulatory systems, involving regions such as the amygdala, thalamus, paracingulate cortex, brainstem, and spinal cord [9, 44, 69, 10, 45]. A second, conceptually distinct possibility is that the source lies in sensory–perceptual processing itself [65]. In this case, variability may arise from altered peripheral input; indeed, chronic pain is associated with changes in spontaneous firing and sensitisation of primary afferents, especially C-fibers, which propagate through the spinal cord to cortical areas [71, 24, 82, 3, 64, 33, 43]. There are also accounts of changes in pain processing regions (e.g. [8]). As our predictive regions span both regulatory and sensory–perceptual components of the pain network, neither mechanism can be identified as the definitive source of variability based on this study. Future research should focus on understanding the origins of variability.

### Variability as marker of pain state

Regardless of its precise origin, two main implications emerge from this study. First, moment-to-moment pain variability provides a simple and interpretable readout of system dynamics, capturing system function over time in an intuitive way: reduced neural variability may reflect a “stiffening” of the system associated with persistently high perceived pain. Second, it offers a tractable means of inferring macro-level changes in the state of the system. Unlike most biomarkers or models that aim to predict absolute pain intensity—a highly subjective measure—variability provides a more objective index that directly links fluctuations in reported pain to the functional state of the system over time. More broadly, compared with recent high-dimensional predictive models that rely on large, multimodal datasets, variability provides a compact descriptor of system state.

Importantly, this measure is straightforward and cost-efficient to implement, requiring only a shift from conventional day-to-day pain reporting to continuous moment-to-moment sampling, alongside targeted fMRI acquisition of predictive regions. Conceptually, it aligns with prior work demonstrating that moment-to-moment within-individual fluctuations can be decoded to track ongoing pain states [46], while extending this framework to show that variability itself captures the system dynamics underlying future clinical trajectories.

Our results indicate that ongoing chronic pain state can be estimated with high accuracy using four longitudinal fMRI sessions, providing a parsimonious representation of the current system state. Building on this, the framework further enables early prediction of future clinical outcomes from a single session acquired at pain onset or during the first 1–2 months post-onset, a stage at which the probability of successful intervention may be highest. This suggests that variability may serve as a clinically actionable marker for early risk stratification without requiring extensive longitudinal sampling. Moreover, the parsimony of the approach is achieved without sacrificing interpretability of the underlying neural substrates, thereby supporting future investigation of targeted interventions.

### Neural correlates of variability follow clinical trajectory

Brain regions associated with moment-to-moment variability largely overlapped with those identified in previous work using the same dataset [7, 62], including alterations in connectivity between insula, subcortical regions, and prefrontal cortex, as well as structural differences in striatal and sensorimotor areas. The convergence of these findings reinforces the view that variability reflects meaningful properties of the pain system rather than noise.

Importantly, although the relative contribution of individual regions varied slightly between pain onset and the early post-onset period, the overall set of predictive regions showed substantial overlap across timepoints. Rather than indicating a sequential reorganisation of the system, this pattern suggests that a stable, distributed network is already established early in the course of the condition, with different nodes contributing differentially depending on sampling and model sensitivity. Core components of this network—including prefrontal, limbic, and sensory–perceptual regions—either remained directly predictive or formed part of overlapping functional systems across both time windows.

Several of the regions identified as predictive have been consistently implicated in pain-related processing across prior neuroimaging studies. These include primary somatosensory cortex, posterior parietal cortex, precuneus, thalamus, amygdala, and multiple prefrontal areas [16, 42, 29, 51, 61, 34, 1, 40, 89, 21, 47, 53, 57, 73, 78, 83, 60]. These regions have been associated with a wide range of processes relevant to pain, including perception, evaluation, salience, and modulation, as well as broader cognitive and affective functions.

Given this functional heterogeneity, the predictive contribution of these regions is unlikely to reflect isolated or specialised roles. Instead, the present results suggest that moment-to-moment variability reflects coordinated activity across distributed neural systems. In this framework, individual regions should be interpreted as components of a larger dynamical network, rather than as discrete functional units selectively encoding specific aspects of pain.

## Limitations

Future work should further investigate the mechanisms underlying variability-related activity and assess whether these regions represent viable targets for intervention. A limitation of the present study is that findings are restricted to back pain and were not validated in independent datasets. Replication across larger and more diverse cohorts will be essential to establish generalisability. Variability may also be combined with existing biomarkers to improve prediction of long-term outcomes.

## Conclusions

Taken together, these findings suggest that moment-to-moment variability in pain provides a window into the dynamical state of pain-regulatory systems. Rather than reflecting noise or epiphenomenal fluctuations, variability appears to index the flexibility of distributed neural networks that govern the evolution of pain over time. The observation that predictive signatures are present at pain onset and persist during early stages indicates that pain chronification is shaped by early-established system dynamics. More broadly, this work highlights variability as a low-dimensional and clinically accessible marker that links neural dynamics to disease progression, opening new avenues for early risk stratification and targeted intervention.

## Methods

### Participants and Experimental Design

Data were obtained from Baliki et al.[7] available through the open access data sharing platform OpenPain [2].The dataset included 120 individuals split into healthy group (24 participants), chronic group (26 participants) and a third group of 70 participants recruited following an episode of subacute back pain (SBP), defined as pain lasting 4–16 weeks with no reported back pain in the preceding year. Participants were followed longitudinally across four visits over one year (Fig 1A): the first within the first month after onset, the second at approximately 1.65±0.10 months, the third at 6.72±0.22 months, and the final at 12.51±0.22 months post-injury. At each visit, participants underwent 3T brain MRI and provided continuous, real-time ratings of spontaneous pain intensity using a finger-spanning device. Ratings were recorded on a Visual Analogue Scale (VAS) ranging from 0 (“no pain”) to 100 (“worst pain imaginable”). The task involved fMRI scanning during spontaneous pain reporting, with a sampling rate of 2.5 seconds, lasting about one hour per session.

Participants were retrospectively classified into two groups based on their average reported pain over the year: persistent pain (SBPp), defined as no meaningful reduction in pain intensity, and recovering pain (SBPr), defined as a progressive decrease corresponding to at least a 20% reduction from baseline. This threshold was adopted to ensure methodological consistency with Baliki et al. [7]. In addition, a group of 24 participants with a formal clinical diagnosis of chronic pain was included as a comparison group.

Demographic information for the participants is summarised in supplementary Table 1. The final sample comprised 51 males and 42 females, with a mean age of 43.5 ± 9.10 years. Participants had an average of 14.2 ± 2.4 years of education. The duration of participation was 8.8 ± 4 months on average. Of the initial cohort, 43 SBPp and 26 SBPr participants were retained for analysis, along with 24 chronic pain patients after data cleaning. Prior work by Baliki et al. [7] reported no significant interaction between group (SBPp vs. SBPr) and analgesic drug use, as quantified using the Methodological Quality Scale (MQS) questionnaire.

Mood and depressive symptoms were assessed using Beck’s Depression Inventory (BDI). At visit 1, mean BDI scores were 6.4 *±* 1.0 for the SBPp group and 6.7 *±* 1.3 for the SBPr group. By visit 4, these values increased to 9.3 *±* 2.1 for SBPp and decreased to 3.8 *±* 0.8 for SBPr.

Short-term affective states were evaluated at both visit 1 and visit 4 using the Positive and Negative Affect Schedule (PANAS). At visit 1, the SBPp group exhibited a positive affect score of 33.4 ± 1.7 and a negative affect score of 22.5 ± 2.6, whereas the SBPr group showed scores of 29.1 ± 2.5 (positive) and 22.7 ± 3.1 (negative). By visit 4, the SBPp group demonstrated scores of 32.5 ± 1.7 (positive) and 20.4 ± 1.7 (negative), while the SBPr group showed 35.4 ± 1.6 (positive) and 14.4 ± 1.1 (negative). Comprehensive details regarding questionnaire responses on drug use, emotional state, depression, mood, and medical conditions are provided in the original study by Baliki et al. [7].

### Participants Inclusion Criteria

Inclusion criteria for each participant required complete neuroimaging data and consistent self-reported pain intensity ratings across sessions. Completeness of pain ratings was defined as continuous engagement with the task throughout each session. Furthermore, at the session level, data from a participant were excluded if they consistently reported pain equal to zero for more than 30% of the session’s duration. The remaining SBPp, SBPr and chronic participants are, respectively, 42, 26 and 24 at onset, 40, 23, 20 at *~*2 months after onset, 43, 24, 21 at *~*7 months after onset, 40, 17,19 at *~*1 year after onset.

### Pain Ratings Analysis

After verifying that each participant had complete imaging data and reported ratings, the validity of each participant’s pain ratings was assessed. A session was considered *invalid* if the participant consistently reported pain equal to 0 for a period equal to or longer than 30% of the session or if the reported ratings ceased before the end of the experiment. In such cases, it was first determined whether an identical session had been conducted for the same participant; if so, data from that session were used. If no replacement data were available, the invalid session was excluded. Following this procedure, the resulting dataset consisted of 43 SBPp, 26 SBPr, and 24 chronic subjects.

Variability in pain rating time series was then analysed for each participant. Given the importance of individual differences in self-reported pain, the coefficient of variation (CV) was employed as a measure of variability:

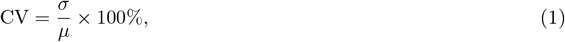

where *σ* is the standard deviation and *µ* is the mean pain rating. The CV is particularly useful as it expresses variability relative to each individual’s mean, thereby accounting for inter-individual differences in reported pain.

### fMRI Preprocessing

To perform the general linear model (GLM) analysis, fMRI data underwent standard cleaning and preprocessing procedures. Data preprocessing was conducted using *fMRIPrep* [23], which included brain extraction, tissue segmentation, spatial normalisation to the MNI152NLin2009cAsym template, and slice-time correction. Subsequent preprocessing and confound regression were performed, following a pipeline consistent with previous work [7] on a subset of the dataset. The applied confounds included motion parameters, white matter and cerebrospinal fluid (WM–CSF) signals, and the global signal. A high-pass temporal filter of 0.006 Hz and spatial smoothing with a 7 mm full-width at half-maximum (FWHM) Gaussian kernel were also applied.

### General Linear Modelling

To identify neural activity linked to pain fluctuations, we employed a two-level voxel-wise General Linear Model (GLM). Raw pain ratings were smoothed to reduce noise and overshoot artifacts prior to modeling. An adaptive low-pass filter was applied, with the cutoff frequency determined automatically from the cumulative power spectrum of each response. A Savitzky–Golay filter was used to compute the second derivative of the spectrum, with the cutoff defined at the point where the derivative approached zero, separating low-frequency pain signals from high-frequency noise.

The smoothed pain ratings, their temporal derivatives, and a movement regressor were then convolved with a canonical Glover hemodynamic response function (HRF) to model the expected BOLD response. This step ensured that the temporal characteristics of the regressors matched the slower dynamics of the BOLD signal, allowing meaningful comparison within the GLM. The movement regressor was derived from abrupt changes in pain ratings, defined as successive differences greater than one unit, capturing rapid shifts but excluding gradual drifts.

Finally, first- and second-level GLMs were fitted to each participant’s whole-brain fMRI data across four sessions, using the three regressors: the temporal derivatives of pain ratings, smoothed pain ratings, and the movement regressor. This analysis yielded voxel-wise z-maps, representing the strength of association between regional BOLD signal fluctuations and the temporal derivatives of the ratings. These z-scores were subsequently used as input features for prediction of clinical outcomes after one year.

### Classifiers

We used Gradient Boosting Machines [28] (GBM; 100 estimators, learning rate 0.1) to classify participants as recovering or persistent. Features consisted of region-of-interest (ROI) values extracted from variability-related BOLD maps. GBM was selected because the relationship between fMRI activity, variability, and clinical outcomes is likely complex and non-linear. GBM is an ensemble of decision trees capable of learning non-linear and interactive effects across features. This is critical in the context of pain chronification, where multiple brain regions interact dynamically [7] and do not simply contribute additively to outcome. GBM also provides measures of feature importance, enabling interpretation of which ROIs most strongly drive prediction. It performs well with relatively small datasets—such as the one we analysed and effectively handles correlated features, which are common in neuroimaging.

Class balance was ensured and the chronic group was treated as comparison group. We studied two main cases: prediction of clinical outcomes after one year using injury onset data, and (ii) prediction of clinical outcomes using data at about 1.6 months from injury onset. The input to the classifier consisted of z-scores representing the relationship between each ROI and variability in reported pain. The output was the class label assigned to each participant. The classifier was evaluated using a stratified 5-fold cross-validation procedure, and performance was measured as the average Receiver Operating Characteristic – Area Under the Curve (ROC AUC) score across folds.

To statistically validate the predictive power of each ROI, a permutation test was performed. An empirical null distribution was generated via whole-dataset permutation, with 1000 iterations in each of the two main cases. Then, an empirical p-value was obtained as:

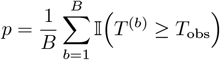

where *T*_obs_ denotes the observed ROC AUC and *T* ^(*b*)^ are the AUC values obtained under label permutation. Only ROIs with *p <*.05 were retained. False Discovery Rate correction was subsequently applied using two approaches: Benjamini–Hochberg (with *α* = 0.05) and Storey’s *Q*-value method (with *λ* = 0.1, *α* = 0.05). Only ROIs identified as significant by both correction methods were considered reliable.

For completeness, we assessed whether our classifier could separate the SBPp and chronic groups at visit 4 based on data from previous visits. For this purpose, we trained our classifier on data from the earliest visit to predict clinical outcomes after *~*1 year and we also trained the classifier on data from *~*1.6 months to predict clinical outcomes after *~*1 year. The results are shown in supplementary figures 1 and 2, respectively.

## Supplementary Material

### Demographic information

Supplementary Table 1 provides detailed demographic information for the participants retained after data cleaning. Only participants included in the final analyses are reported, and for each participant only visits meeting completeness and data-quality criteria were retained. The “visit” column indicates the visits attended by each participant that were considered valid and included in the analyses, as described in the Methods section. Additional demographic variables, including gender, age, ethnicity, and years of education, are also reported. Further details regarding PANAS and MQC questionnaire measures assessing mood, depression, and related conditions can be found in the original study by Baliki et al. [9].

**Table 1.**
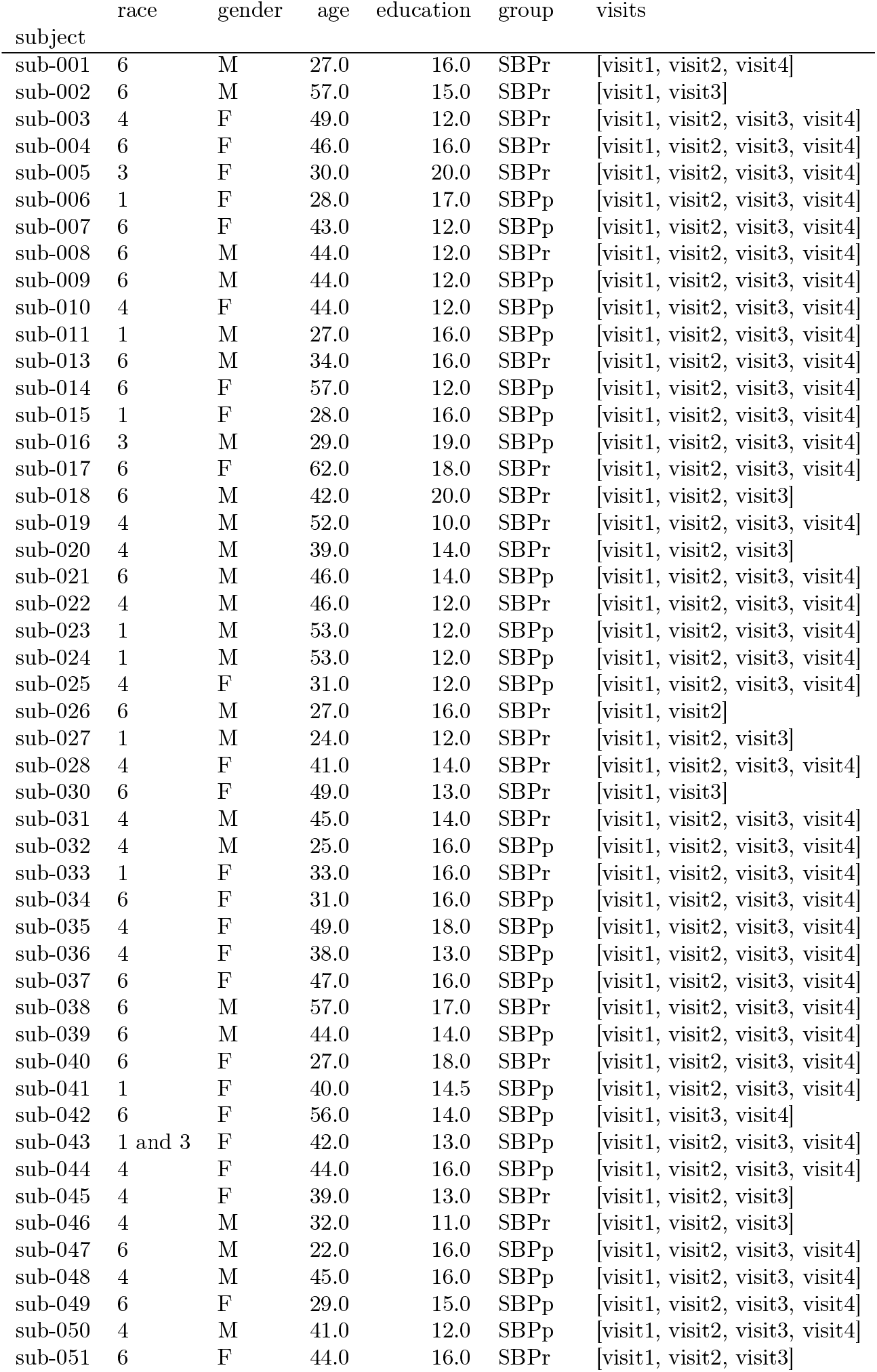
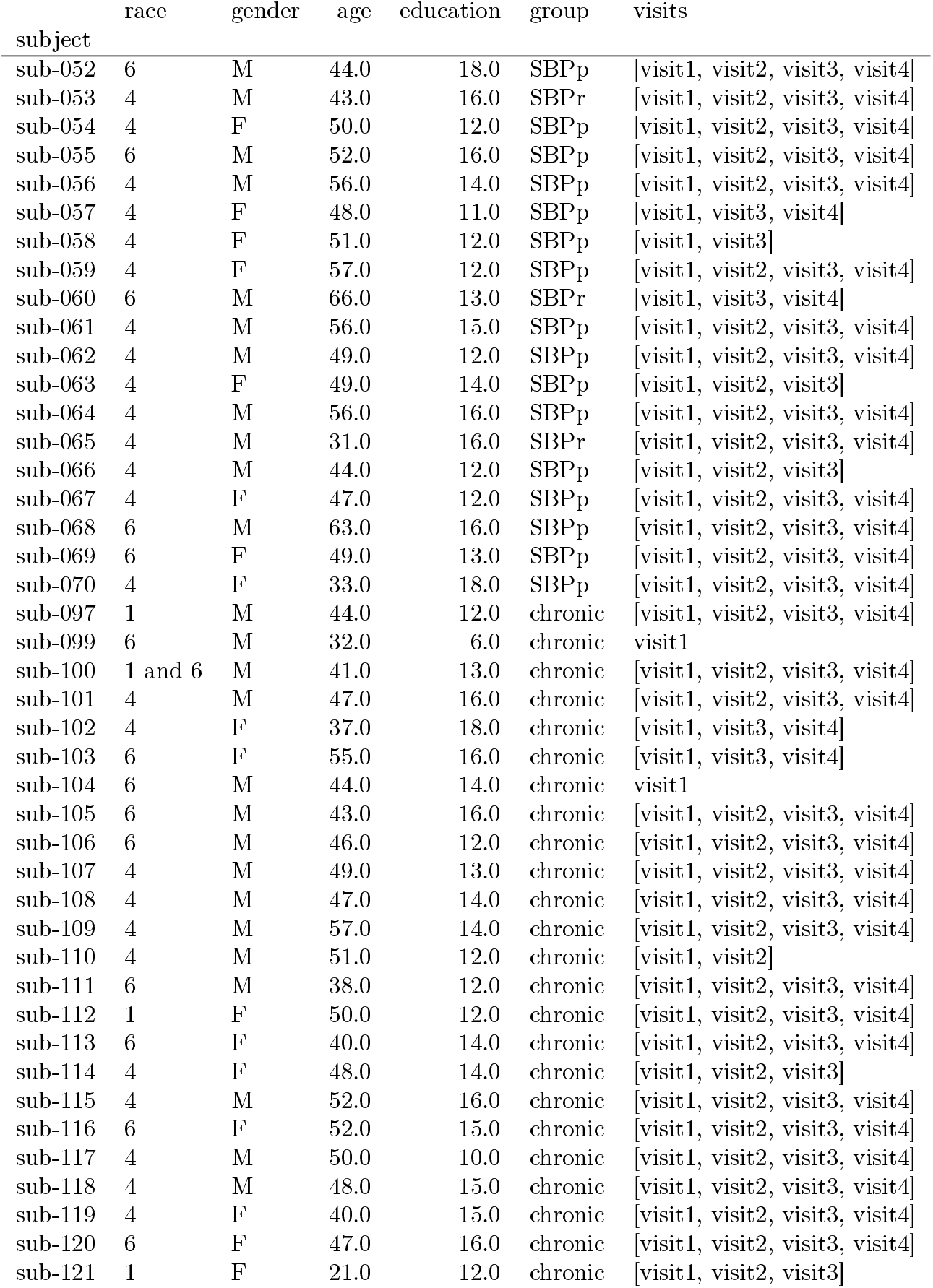
Demographic characteristics of the participants selected for this study, drawn from the original dataset reported by Baliki et al. The columns are defined as follows: *race* denotes the participants’ ethnicity, which is reported using a coded scheme, as shown in supplementary Table 1. These codes correspond to the labels defined in [7]; *education* indicates years of formal education; *duration* duration of participation in months; *group* specifies the classification assigned to each participant (see Methods); and *visits* lists the visits attended and retained as valid after data cleaning.

|  | race | gender | age | education | group | visits |
| --- | --- | --- | --- | --- | --- | --- |
| subject |  |  |  |  |  |  |
| sub-001 | 6 | M | 27.0 | 16.0 | SBPr | [visit1, visit2, visit4] |
| sub-002 | 6 | M | 57.0 | 15.0 | SBPr | [visit1, visit3] |
| sub-003 | 4 | F | 49.0 | 12.0 | SBPr | [visit1, visit2, visit3, visit4] |
| sub-004 | 6 | F | 46.0 | 16.0 | SBPr | [visit1, visit2, visit3, visit4] |
| sub-005 | 3 | F | 30.0 | 20.0 | SBPr | [visit1, visit2, visit3, visit4] |
| sub-006 | 1 | F | 28.0 | 17.0 | SBPp | [visit1, visit2, visit3, visit4] |
| sub-007 | 6 | F | 43.0 | 12.0 | SBPp | [visit1, visit2, visit3, visit4] |
| sub-008 | 6 | M | 44.0 | 12.0 | SBPr | [visit1, visit2, visit3, visit4] |
| sub-009 | 6 | M | 44.0 | 12.0 | SBPp | [visit1, visit2, visit3, visit4] |
| sub-010 | 4 | F | 44.0 | 12.0 | SBPp | [visit1, visit2, visit3, visit4] |
| sub-011 | 1 | M | 27.0 | 16.0 | SBPp | [visit1, visit2, visit3, visit4] |
| sub-013 | 6 | M | 34.0 | 16.0 | SBPr | [visit1, visit2, visit3, visit4] |
| sub-014 | 6 | F | 57.0 | 12.0 | SBPp | [visit1, visit2, visit3, visit4] |
| sub-015 | 1 | F | 28.0 | 16.0 | SBPp | [visit1, visit2, visit3, visit4] |
| sub-016 | 3 | M | 29.0 | 19.0 | SBPp | [visit1, visit2, visit3, visit4] |
| sub-017 | 6 | F | 62.0 | 18.0 | SBPr | [visit1, visit2, visit3, visit4] |
| sub-018 | 6 | M | 42.0 | 20.0 | SBPr | [visit1, visit2, visit3] |
| sub-019 | 4 | M | 52.0 | 10.0 | SBPr | [visit1, visit2, visit3, visit4] |
| sub-020 | 4 | M | 39.0 | 14.0 | SBPr | [visit1, visit2, visit3] |
| sub-021 | 6 | M | 46.0 | 14.0 | SBPp | [visit1, visit2, visit3, visit4] |
| sub-022 | 4 | M | 46.0 | 12.0 | SBPr | [visit1, visit2, visit3, visit4] |
| sub-023 | 1 | M | 53.0 | 12.0 | SBPp | [visit1, visit2, visit3, visit4] |
| sub-024 | 1 | M | 53.0 | 12.0 | SBPp | [visit1, visit2, visit3, visit4] |
| sub-025 | 4 | F | 31.0 | 12.0 | SBPp | [visit1, visit2, visit3, visit4] |
| sub-026 | 6 | M | 27.0 | 16.0 | SBPr | [visit1, visit2] |
| sub-027 | 1 | M | 24.0 | 12.0 | SBPr | [visit1, visit2, visit3] |
| sub-028 | 4 | F | 41.0 | 14.0 | SBPr | [visit1, visit2, visit3, visit4] |
| sub-030 | 6 | F | 49.0 | 13.0 | SBPr | [visit1, visit3] |
| sub-031 | 4 | M | 45.0 | 14.0 | SBPr | [visit1, visit2, visit3, visit4] |
| sub-032 | 4 | M | 25.0 | 16.0 | SBPp | [visit1, visit2, visit3, visit4] |
| sub-033 | 1 | F | 33.0 | 16.0 | SBPr | [visit1, visit2, visit3, visit4] |
| sub-034 | 6 | F | 31.0 | 16.0 | SBPp | [visit1, visit2, visit3, visit4] |
| sub-035 | 4 | F | 49.0 | 18.0 | SBPp | [visit1, visit2, visit3, visit4] |
| sub-036 | 4 | F | 38.0 | 13.0 | SBPp | [visit1, visit2, visit3, visit4] |
| sub-037 | 6 | F | 47.0 | 16.0 | SBPp | [visit1, visit2, visit3, visit4] |
| sub-038 | 6 | M | 57.0 | 17.0 | SBPr | [visit1, visit2, visit3, visit4] |
| sub-039 | 6 | M | 44.0 | 14.0 | SBPp | [visit1, visit2, visit3, visit4] |
| sub-040 | 6 | F | 27.0 | 18.0 | SBPr | [visit1, visit2, visit3, visit4] |
| sub-041 | 1 | F | 40.0 | 14.5 | SBPp | [visit1, visit2, visit3, visit4] |
| sub-042 | 6 | F | 56.0 | 14.0 | SBPp | [visit1, visit3, visit4] |
| sub-043 | 1 and 3 | F | 42.0 | 13.0 | SBPp | [visit1, visit2, visit3, visit4] |
| sub-044 | 4 | F | 44.0 | 16.0 | SBPp | [visit1, visit2, visit3, visit4] |
| sub-045 | 4 | F | 39.0 | 13.0 | SBPr | [visit1, visit2, visit3] |
| sub-046 | 4 | M | 32.0 | 11.0 | SBPr | [visit1, visit2, visit3] |
| sub-047 | 6 | M | 22.0 | 16.0 | SBPp | [visit1, visit2, visit3, visit4] |
| sub-048 | 4 | M | 45.0 | 16.0 | SBPp | [visit1, visit2, visit3, visit4] |
| sub-049 | 6 | F | 29.0 | 15.0 | SBPp | [visit1, visit2, visit3, visit4] |
| sub-050 | 4 | M | 41.0 | 12.0 | SBPp | [visit1, visit2, visit3, visit4] |
| sub-051 | 6 | F | 44.0 | 16.0 | SBPr | [visit1, visit2, visit3] |
| subject |  |  |  |  |  |  |
| sub-052 | 6 | M | 44.0 | 18.0 | SBPp | [visit1, visit2, visit3, visit4] |
| sub-053 | 4 | M | 43.0 | 16.0 | SBPr | [visit1, visit2, visit3, visit4] |
| sub-054 | 4 | F | 50.0 | 12.0 | SBPp | [visit1, visit2, visit3, visit4] |
| sub-055 | 6 | M | 52.0 | 16.0 | SBPp | [visit1, visit2, visit3, visit4] |
| sub-056 | 4 | M | 56.0 | 14.0 | SBPp | [visit1, visit2, visit3, visit4] |
| sub-057 | 4 | F | 48.0 | 11.0 | SBPp | [visit1, visit3, visit4] |
| sub-058 | 4 | F | 51.0 | 12.0 | SBPp | [visit1, visit3] |
| sub-059 | 4 | F | 57.0 | 12.0 | SBPp | [visit1, visit2, visit3, visit4] |
| sub-060 | 6 | M | 66.0 | 13.0 | SBPr | [visit1, visit3, visit4] |
| sub-061 | 4 | M | 56.0 | 15.0 | SBPp | [visit1, visit2, visit3, visit4] |
| sub-062 | 4 | M | 49.0 | 12.0 | SBPp | [visit1, visit2, visit3, visit4] |
| sub-063 | 4 | F | 49.0 | 14.0 | SBPp | [visit1, visit2, visit3] |
| sub-064 | 4 | M | 56.0 | 16.0 | SBPp | [visit1, visit2, visit3, visit4] |
| sub-065 | 4 | M | 31.0 | 16.0 | SBPr | [visit1, visit2, visit3, visit4] |
| sub-066 | 4 | M | 44.0 | 12.0 | SBPp | [visit1, visit2, visit3] |
| sub-067 | 4 | F | 47.0 | 12.0 | SBPp | [visit1, visit2, visit3, visit4] |
| sub-068 | 6 | M | 63.0 | 16.0 | SBPp | [visit1, visit2, visit3, visit4] |
| sub-069 | 6 | F | 49.0 | 13.0 | SBPp | [visit1, visit2, visit3, visit4] |
| sub-070 | 4 | F | 33.0 | 18.0 | SBPp | [visit1, visit2, visit3, visit4] |
| sub-097 | 1 | M | 44.0 | 12.0 | chronic | [visit1, visit2, visit3, visit4] |
| sub-099 | 6 | M | 32.0 | 6.0 | chronic | visit1 |
| sub-100 | 1 and 6 | M | 41.0 | 13.0 | chronic | [visit1, visit2, visit3, visit4] |
| sub-101 | 4 | M | 47.0 | 16.0 | chronic | [visit1, visit2, visit3, visit4] |
| sub-102 | 4 | F | 37.0 | 18.0 | chronic | [visit1, visit3, visit4] |
| sub-103 | 6 | F | 55.0 | 16.0 | chronic | [visit1, visit3, visit4] |
| sub-104 | 6 | M | 44.0 | 14.0 | chronic | visit1 |
| sub-105 | 6 | M | 43.0 | 16.0 | chronic | [visit1, visit2, visit3, visit4] |
| sub-106 | 6 | M | 46.0 | 12.0 | chronic | [visit1, visit2, visit3, visit4] |
| sub-107 | 4 | M | 49.0 | 13.0 | chronic | [visit1, visit2, visit3, visit4] |
| sub-108 | 4 | M | 47.0 | 14.0 | chronic | [visit1, visit2, visit3, visit4] |
| sub-109 | 4 | M | 57.0 | 14.0 | chronic | [visit1, visit2, visit3, visit4] |
| sub-110 | 4 | M | 51.0 | 12.0 | chronic | [visit1, visit2] |
| sub-111 | 6 | M | 38.0 | 12.0 | chronic | [visit1, visit2, visit3, visit4] |
| sub-112 | 1 | F | 50.0 | 12.0 | chronic | [visit1, visit2, visit3, visit4] |
| sub-113 | 6 | F | 40.0 | 14.0 | chronic | [visit1, visit2, visit3, visit4] |
| sub-114 | 4 | F | 48.0 | 14.0 | chronic | [visit1, visit2, visit3] |
| sub-115 | 4 | M | 52.0 | 16.0 | chronic | [visit1, visit2, visit3, visit4] |
| sub-116 | 6 | F | 52.0 | 15.0 | chronic | [visit1, visit2, visit3, visit4] |
| sub-117 | 4 | M | 50.0 | 10.0 | chronic | [visit1, visit2, visit3, visit4] |
| sub-118 | 4 | M | 48.0 | 15.0 | chronic | [visit1, visit2, visit3, visit4] |
| sub-119 | 4 | F | 40.0 | 15.0 | chronic | [visit1, visit2, visit3, visit4] |
| sub-120 | 6 | F | 47.0 | 16.0 | chronic | [visit1, visit2, visit3, visit4] |
| sub-121 | 1 | F | 21.0 | 12.0 | chronic | [visit1, visit2, visit3] |

## Supplementary analyses

For completeness, we performed additional analyses considering only the SBPp and chronic groups, testing whether neural correlates of moment-to-moment pain variability could distinguish between individuals who later developed persistent pain and those already exhibiting chronic pain. Classification models were trained using data acquired either at pain onset or during the early post-onset period (~1.6 months after onset), with prediction targets defined at one year. Results are reported in Figures 1 and 2.

**Figure 1.**
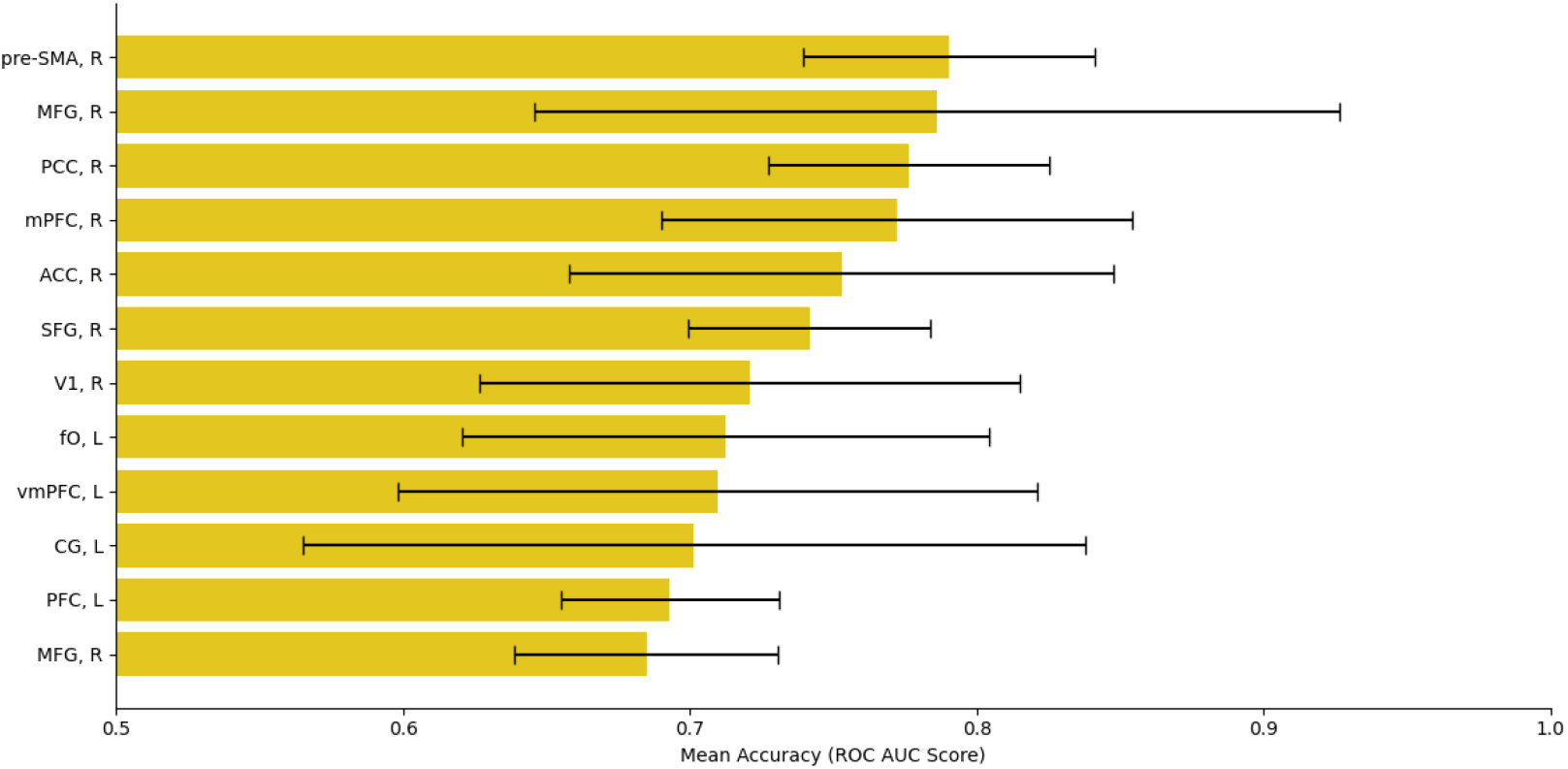
Top 12 ROIs contributing to classification performance using a Gradient Boosting Model (GBM) classifier trained on data from the earliest visit to predict clinical outcomes at one year. ROI labels correspond to the following anatomical regions: *pre-SMA, R*: Right Pre-Supplementary Motor Area; *MFG, R*: Right Middle Frontal Gyrus; *PCC, R*: Right Posterior Cingulate Cortex; *mPFC, R*: Right medial Prefrontal Cortex; *ACC, R*: Right Anterior Cingulate Cortex; *SFG, R*: Right Superior Frontal Gyrus; *V1, R*: Right Primary Visual Cortex; *fO, L*: Left Frontal Operculum; *vmPFC, L*: Left ventro medial Prefrontal Cortex; *CG, L*: Left Cingulate Gyrus; *PFC, L*: Left Prefrontal Cortex; *MFG, R*: Right Middle Frontal Gyrus.

**Figure 2.**
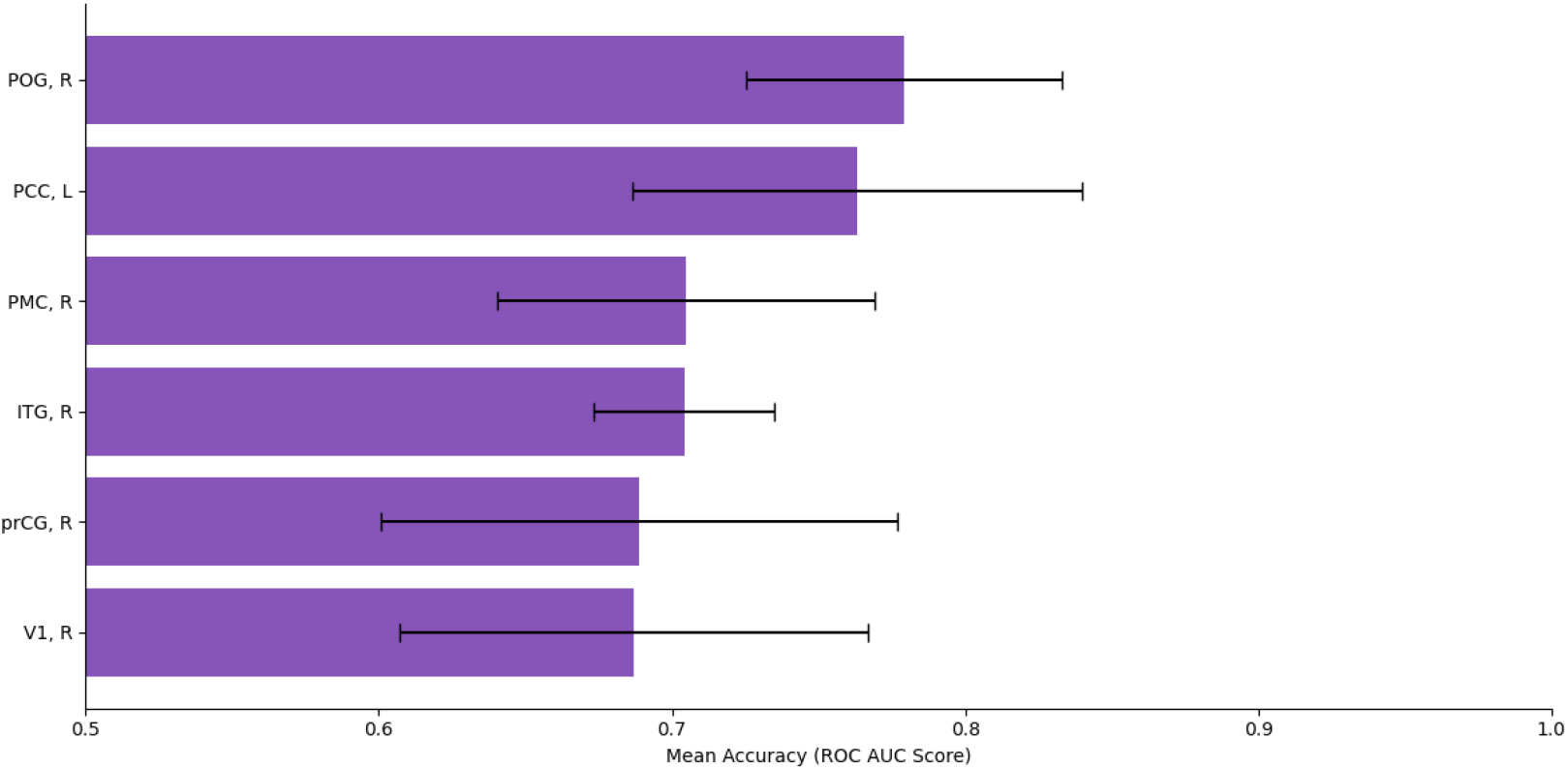
Top 12 ROIs contributing to classification performance using a Gradient Boosting Model (GBM) classifier trained on data from the second visit ~1.6 months after onset, to predict clinical outcomes at one year. ROI labels correspond to the following anatomical regions: *POG, R*: Right Postcentral Gyrus; *PCC, L*: Left Posterior Cingulate Gyrus; *PMC, R*: Right Premotor Cortex; *ITG, R*: Right Inferior Temporal Gyrus; *prCG, R*: Right Precentral Gyrus; *V1, R*: Right Primary Visual Cortex.

Overall, classification performance was reduced relative to analyses including the recovering group, indicating greater similarity between SBPp and chronic participants. Nevertheless, predictive information remained detectable at the earliest visit, suggesting that neural signatures associated with chronic pain are already established shortly after onset. By the second visit, the classifier showed increasing difficulty distinguishing SBPp from chronic participants, consistent with the interpretation that the SBPp group progressively converges toward a chronic pain-like state over time.

**Table 2.** MNI coordinates of the top 12 regions whose activity associated with moment-to-moment pain variability in SBPp and chronic participants at pain onset most strongly predicted clinical outcome at one year. Here, SBPr participants are excluded. Regions are ranked based on their predictive contribution in the classification model. Values represent mean ROC AUC across 5-fold cross-validation.

| Region | Coordinates (MNI152) |
| --- | --- |
| Right Pre-Supplementary Motor Area (Pre-SMA) | [−4.69, −28.88, 26.88] |
| Right Middle Frontal Gyrus (MFG) | [25.98, 7.09, 57.66] |
| Right Posterior Cingulate Cortex (PCC) | [12.85, −54.55, 15.02] |
| Right medial Prefrontal Cortex (MPFC) | [29.73, 48.13, 27.12] |
| Right Anterior Cingulate Cortex (ACC) | [7.89, 41.93, 4.03] |
| Right Superior Frontal Gyrus (SFG) | [7.85, 25.58, 54.74] |
| Right Primary Visual Cortex (V1) | [42.25, −79.83, 9.7] |
| Left Frontal Operculum (fO) | [−36.92, −20.96, 15.28] |
| Left ventro medial Prefrontal Cortex (vmPFC) | [−27.54, 58.03, 8.04] |
| Left Cingulate Gyrus (CG) | [−4.69, −28.88, 26.88] |
| Left Prefrontal Cortex (PFC) | [−13.25, 62.56, −5.7] |
| Right Middle Frontal Gyrus (MFG) | [40.62, 32.95, 37.24] |

**Table 3.** MNI coordinates of the top 12 regions whose activity associated with moment-to-moment pain variability in SBPp and chronic participants at ~1.6 months after onset most strongly predicted clinical outcome at one year. Here, SBPr participants are excluded. Regions are ranked based on their predictive contribution in the classification model. Values represent mean ROC AUC across 5-fold cross-validation.

| Region | Coordinates (MNI152) |
| --- | --- |
| Right Postcentral Gyrus (POG) | [50.9, -22.41, 51.84] |
| Left Posterior Cingulate Gyrus (PCC, L) | [-4.69, -28.88, 26.88] |
| Right Premotor Cortex (PMC, R) | [42.13, 14.28, 48.97] |
| Right Inferior Temporal Gyrus (ITG) | [62.59, -41.91, -11.43] |
| Right Precentral Gyrus (prCG) | [59.01, 0.55, 10.93] |
| Right Primary Visual Cortex (V1) | [26.26, -61.28, 57.97] |

## Acknowledgments

FM is funded by a UKRI Medical Research Council Career Development Award (MR/T010614/1), a UKRI Advanced Pain Discovery Platform grant (MR/ W027593/1), and an UKRI Engineering and Physical Sciences Research Council +Medical Research Council Programme Grant (UKRI1970). GP is also funded by MR/W027593/1. CA is funded by an EPSRC DTP (EP/W524633/1).

